# Mechanistic view and genetic control of DNA recombination during meiosis

**DOI:** 10.1101/169870

**Authors:** Marie-Claude Marsolier-Kergoat, Md Muntaz Khan, Jonathan Schott, Xuan Zhu, Bertrand Llorente

## Abstract

Meiotic recombination is essential for fertility and allelic shuffling. Canonical recombination models fail to capture the observed complexity of meiotic recombinants. Here we revisit these models by analyzing meiotic heteroduplex DNA tracts genome-wide in combination with meiotic DNA double-strand break (DSB) locations. We provide unprecedented support to the synthesis-dependent strand annealing model and establish estimates of its associated template switching frequency and polymerase processivity. We show that resolution of double Holliday junctions (dHJs) is biased toward cleavage of the pair of strands containing newly synthesized DNA near the junctions. The suspected dHJ resolvase Mlh1-3 as well as Mlh1-2, Exo1 and Sgs1 promote asymmetric positioning of crossover intermediates relative to the initiating DSB and bidirectional conversions. Finally, we show that crossover-biased dHJ resolution depends on Mlh1-3, Exo1, Msh5 and to a lesser extent on Sgs1. These properties are likely conserved in eukaryotes containing the ZMM proteins, which includes mammals.

## INTRODUCTION

Decades of studies at a few loci in different organisms led to a global view of meiotic recombination (Figure 1). In the representative *Saccharomyces cerevisiae* yeast model, it starts by the formation of DNA double-strand breaks (DSBs) catalyzed by the Spo11 topoisomerase II-like protein^1, 2^. Second, DNA ends are processed by the Mre11-Rad50-Xrs2 complex and Sae2 to release Spo11 from the ends, which allows Exo1 to further degrade the 5’ ends and produce on average ca. 800 bp-long 3’ single-stranded DNA tails^3-7^. Third, a DNA tail invades a homologous template to form a D-loop structure comprising a heteroduplex DNA (hDNA) plus a 3’ end used to prime DNA repair synthesis.

**Figure 1:**
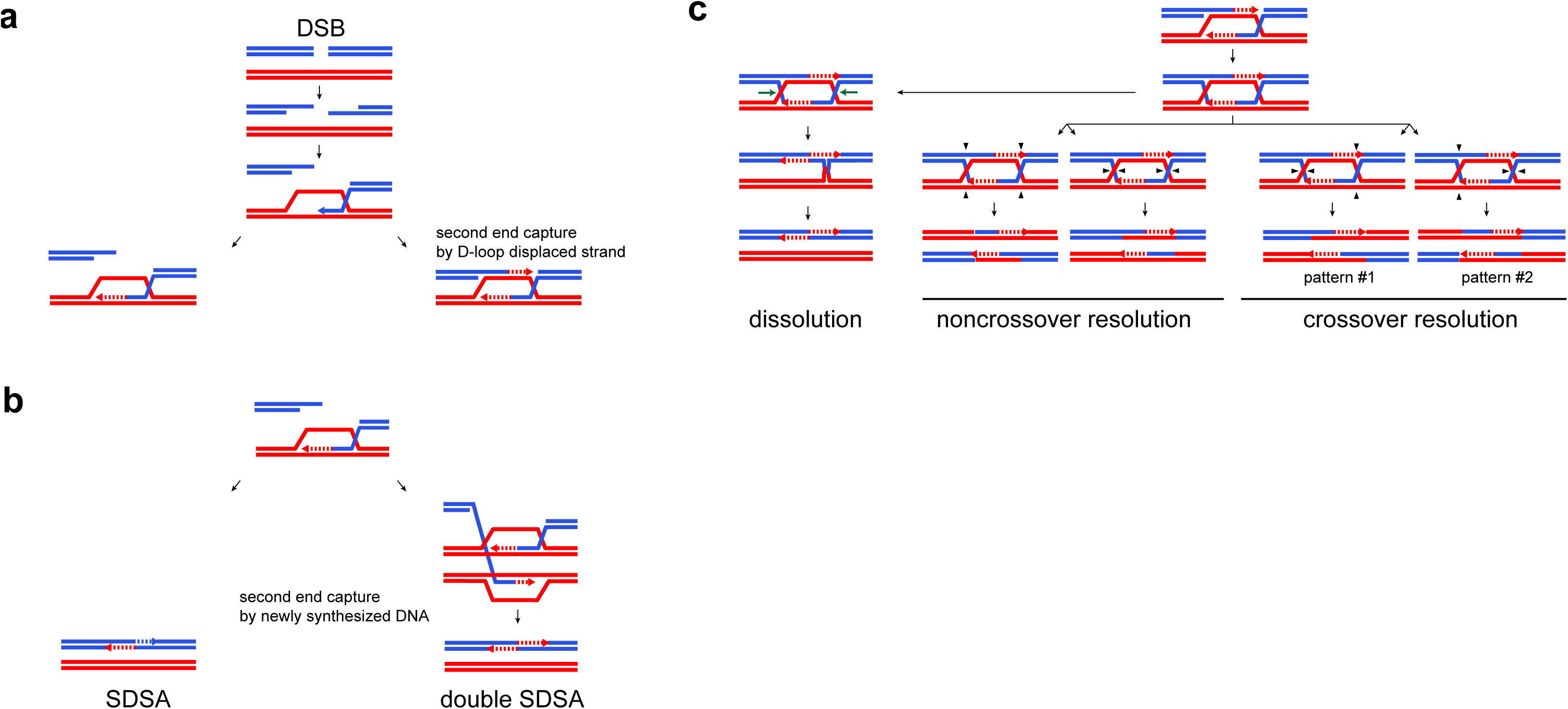
Canonical recombination pathways in the absence of mismatch repair. **a**. Meiotic DSB processing. Homologous double-stranded DNA molecules are represented by pairs of parallel blue and red lines. The arrowheads indicate the 5’ to 3’ polarity when needed. Dotted lines represent newly synthesized DNA. The left product with no capture of the second end by the displaced strand of the D-loop is channeled in the SDSA/double SDSA pathway (panel B). The right product is channeled in the double Holliday junction processing pathway (panel C). **b**. SDSA/double SDSA pathways. **c**. Processing of dHJs. Horizontal green arrows indicate the direction of HJ branch migration during dissolution. Plain black arrowheads represent the nick locations required for HJ resolution. There are two resolution configurations for both noncrossovers and crossovers. The two end products are identical for noncrossovers, but are distinct for crossovers. In pattern #1, the hDNA tracts are in continuity with both parental duplexes, while they are in discontinuity in pattern #2.

Next, under the synthesis-dependent strand annealing (SDSA) model, the resected second end anneals to the newly synthesized DNA on the first end after D-loop disruption to yield exclusively noncrossovers^8–11^. The first hDNA formed is transient but the second hDNA (newly synthesized DNA annealed to the second end) may be present in the final noncrossover just 3’ of the initiating DSB (Figure 1B and 2B).

**Figure 2:**
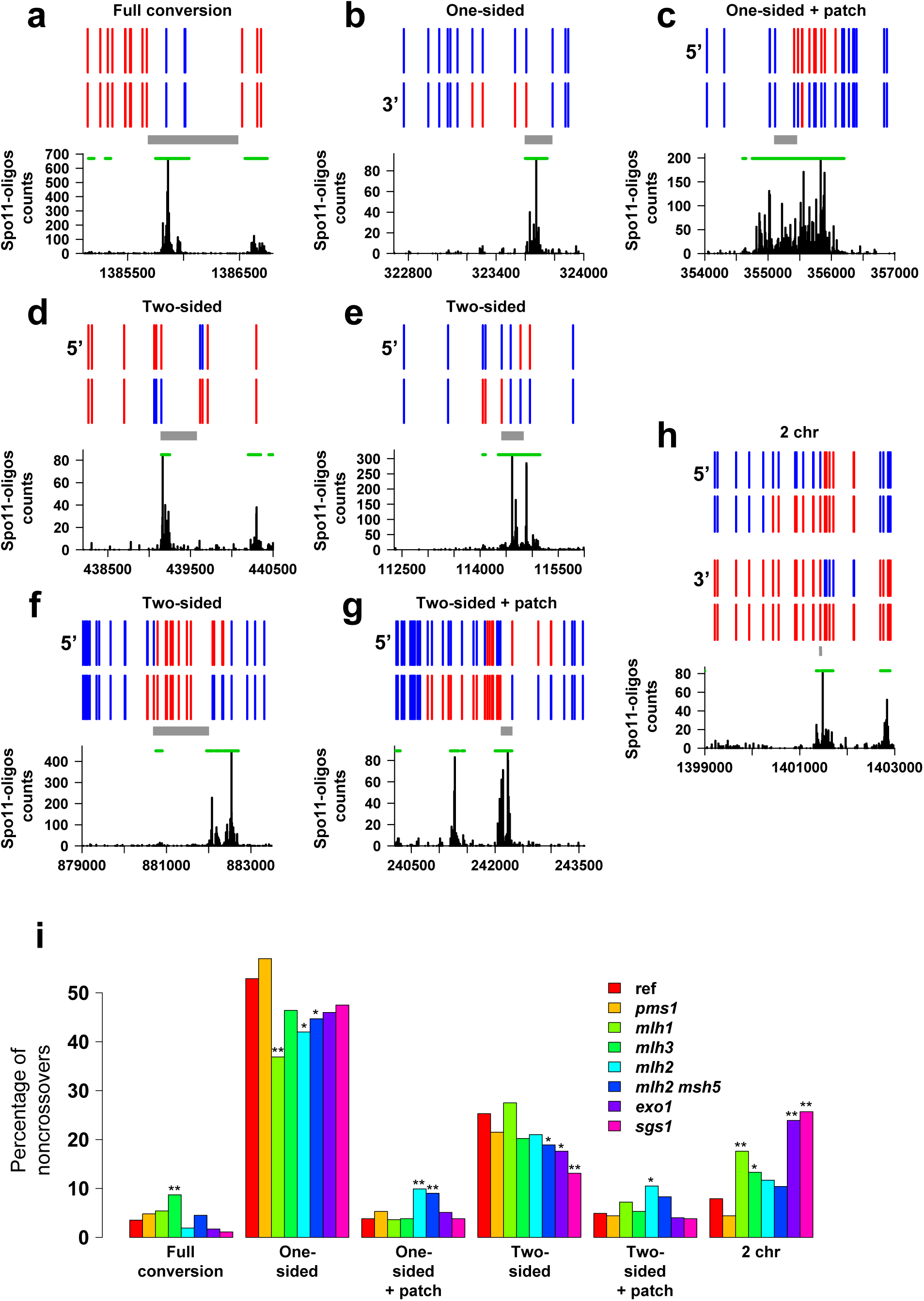
Examples and categorization of the noncrossovers detected in the *msh2A* octads. (**a**-**h**) A representative set of noncrossovers involving one chromatid or two non-sister chromatids. In the top part of each figure, vertical bars represent SNPs locations, with the S288C and SK1 alleles as red and blue lines, respectively. Each horizontal series of vertical bars is a DNA strand. One (**a**-**g**) or two (**h**) chromatids are shown. The green lines represent the dsb95 segments. The grey rectangles correspond to the expected locations of the initiating DSBs according to the current recombination models (see Figure 1). The bottom part of each figure shows the counts of immunoprecipitated Spo11-FLAG oligos for each position, using S288C coordinates. **a**. Noncrossover with a single full-conversion tract that is compatible with gap repair. **b**. One-sided noncrossover with a half-conversion tract only. **c**. Onesided noncrossover with a half-conversion tract and an internal patch of full conversion. **d**. Two-sided noncrossover with two half-conversion tracts (trans hDNA) affecting the same chromatid. **e**. Two-sided noncrossover with two half-conversion tracts affecting the same chromatid and separated by a restoration tract. **f**. Two-sided noncrossover with two half-conversion tracts affecting the same chromatid and separated by a full-conversion tract. **g**. Two-sided noncrossover with two halfconversion tracts affecting the same chromatid and an internal patch of full conversion. **h**. Two-sided noncrossovers affecting two non-sister chromatids. **j**. Noncrossovers categories. Noncrossovers were classified according to their strand transfer pattern illustrated in panels **a**-**h** and the percentage of each category over the total number of noncrossovers is shown. The stars indicate when the abundance of the noncrossover category in a mutant is significantly different from the reference *msh2*Δ strain (one star, *p* < 0.05; two stars, *p* < 0.01, Fisher’s exact test).

Alternatively, under the canonical DSB repair model, the resected second end anneals to the displaced strand of the D-loop, yielding an intermediate with two Holliday junctions (dHJ). Depending on whether two or four DNA strands are cleaved, dHJ resolution yields either a noncrossover or a crossover, respectively (Figure 1C)^4, 12^.

While crossovers result from dHJ resolution, meiotic noncrossovers are thought to mostly result from SDSA and dHJ dissolution by isomerization through branch migration yielding trans hDNA tracts on the same chromatid (Figure 1B and 2D)^13–15^. In many organisms, including *S. cerevisiae*, plants, and mammals, crossovers form through two distinct pathways. The major pathway relies on the integrity of the synaptonemal complex and the resulting crossovers show interference, *i.e.* they are more widely and evenly spaced than expected by chance. In yeast, this ZMM pathway depends on Zip1, Zip2, Zip3, Zip4, Spo16, the Mer3 helicase and the heterodimer Msh4-Msh5 thought to protect recombination intermediates from Sgs1-mediated dissolution^16, 17^. The heterodimeric endonuclease Mlh1-Mlh3 in combination with Sgs1 and Exo1 are thought to constitute the resolution machinery of the ZMM pathway^18–21^. The minor crossover pathway is independent of the synaptonemal complex and the resulting crossovers do not show interference^22–24^. It relies on the partially redundant structure-specific nucleases Mus81-Mms4, Yen1 and Slx1-Slx4^18, 22, 25, 26^. Interestingly, dHJ resolution is biased toward crossover in the ZMM pathway, but apparently not in the structure specific nulcease pathway(s)^18, 27, 28, 29, 30^. The mechanism for this bias is not known.

The hDNA tracts disposition in recombinants is key to understanding recombination mechanisms. To be studied, they require either inactivation of mismatch repair^31^ or poorly repairable substrates^32^. Under the canonical DSB repair model, dHJ resolution yields hDNA tracts on both sides of the initiating DSB. However, hDNA tracts were instead frequently observed on one side only^13–15, 33–36^ (Figure 3C). These one-sided events could reflect either a structural asymmetry of the hDNA tracts formed early in the recombination reaction, or that some hDNA tracts are transient.

**Figure 3:**
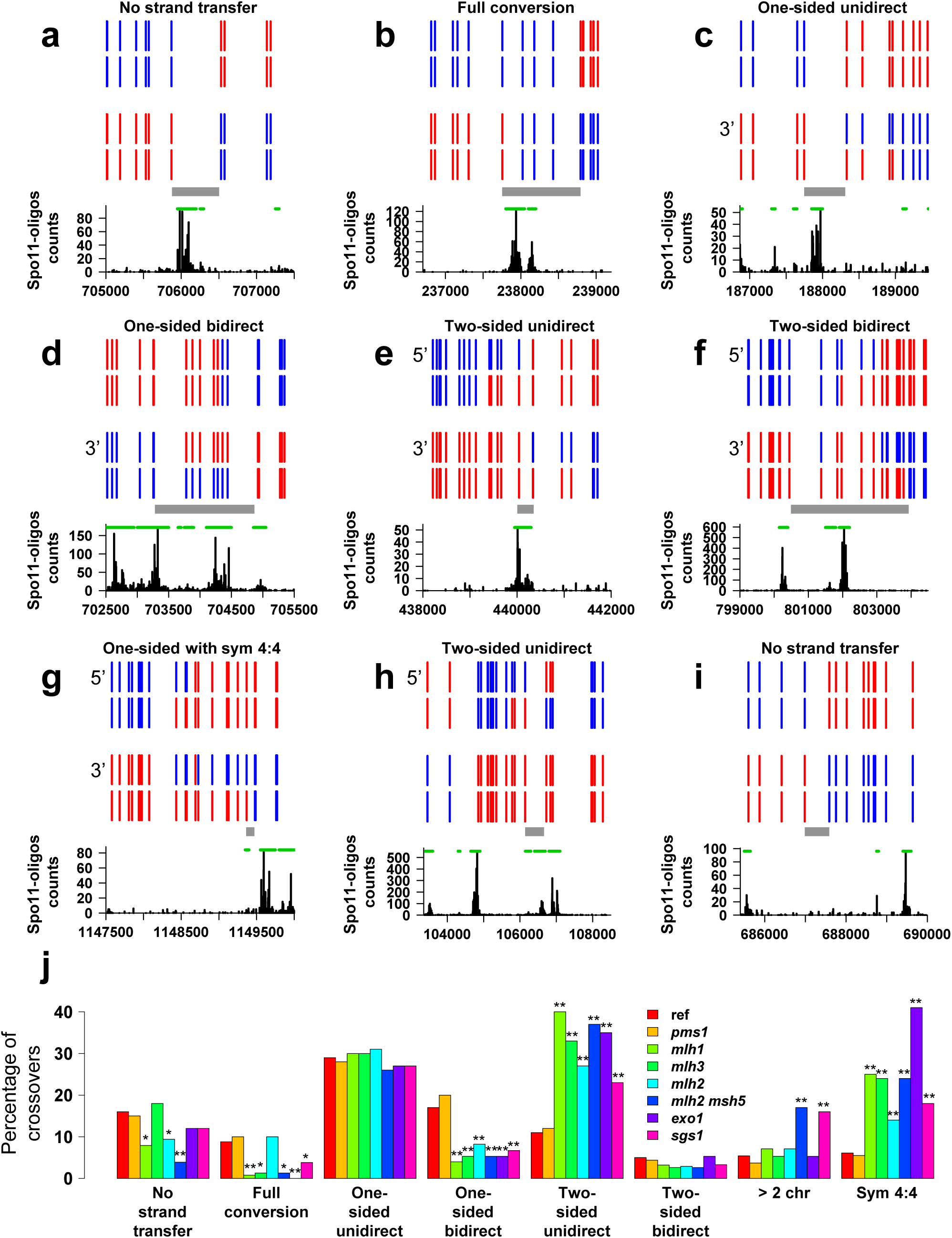
Examples and categorization of the crossovers detected in the *msh2*Δ octads. (**a**-**i**) A representative set of crossovers involving two non-sister chromatids are schematized as in Figure 2. **a**. Crossover belonging to the category “no strand transfer” in Figure 3J. **b**. Crossover with a single full-conversion tract. **c**. One-sided, unidirectional crossover composed of a single half-conversion tract. **d**. One-sided, bidirectional crossover. **e**. Two-sided, unidirectional crossover with trans hDNA tracts on the two recombining chromatids. **f**. Two-sided, bidirectional crossover. **g**. Crossover with a terminal tract of symmetrical hDNA possibly originating from HJ branch migration. **h**. Two-sided, unidirectional crossover with trans hDNA tracts on one chromatid only. **i**. Crossover belonging to the category “no strand transfer” without any dsb95 segment located at the position expected for the causal DSB. **j**. Crossover categories. Crossovers were classified according to their strand transfer pattern (see panels **a**-**i**) and the percentage of each category over the total number of crossovers is shown. The first seven categories are mutually exclusive. The stars indicate when the abundance of the crossover category in a mutant is significantly different from the reference *msh2*Δ strain (one star, *p* < 0.05; two stars, *p* < 0.01, Fisher’s exact test).

Another prediction of the canonical model is that crossover-associated hDNA should occur equally frequently in either of two configurations (patterns 1 and 2 in Figure 1C), because both HJs should have equal likelihood of being resolved as a crossover. However, a prevalence of crossovers occurring at the junction inferred to result from the first strand invasion was observed, albeit to different extents among the few loci analyzed^33, 34, 37^. As this configuration requires cleavage of strands proximal to the newly synthesized DNA tracts, it suggests that these tracts somehow bias the orientation of HJ resolution.

Finally, the gain of genetic material at a recombination site is expected to be unidirectional, from the intact donor to the broken recipient. However, bidirectional exchanges of genetic information have been reported in many genetic studies (Figure 3D and E), including those designed to follow recombination events arising from a unique DSB hotspot^34^. These were considered to be the result of multiple independent Spo11-DSBs and were disregarded.

We recently developed a strategy to analyze meiotic hDNA genome-wide from a S288C × SK1 hybrid lacking the Msh2 mismatch repair component (Figure S1)^15^. Despite new insights provided, a major limitation was the lack of knowledge of the location of the initiating DSBs, as for any other genome-wide study of recombination so far. Therefore, here, we explore the discrepancies between canonical models and *in vivo* recombination patterns by combining maps of genome-wide Spo11-DSB locations with strand-transfer patterns in a mismatch repair-defective reference strain and different recombination mutants. The results provide unprecedented detail about global patterns and genetic control of recombination and present a framework for revising and refining mechanistic recombination models.

## RESULTS

### Matching recombination events to initiating DSBs

Through sequencing Spo11-bound oligonucleotides (Spo11-oligos), we determined the meiotic DSB map for the S288C × SK1 hybrid, which closely resembles that of the SK1 × SK1 homozygote^38, 39^. In parallel we analyzed the progeny of four octads derived from a diploid *msh2*Δ strain and inferred the most parsimonious scenarios for the strand transfers characterizing recombination events^40^. Figures 2A-H and 3A-I show representative examples of the 367 noncrossovers and 448 crossovers observed and listed in Table S1. In order to give a comprehensive and synthetic description of these events, we coded them with series of colored segments (Figure S2-5).

The Spo11-oligo map remarkably fits with the expected positions of initiating DSBs for the recombination events (Figures 2A-H and 3A-H). To quantify the goodness of this fit, the genome was divided into 50-bp segments and only the segments harboring ≥ 16 associated Spo11-oligos were considered as DSB-containing regions. Collectively, these DSB regions contain 95% of all Spo11-oligos and cover 15% of the genome. We reasoned that for any set of recombination events, 95% of the initiating DSBs should be located within these DSB regions (also called “dsb95 segments”) and we quantified the discrepancies between the expected locations of DSBs inferred from the current recombination models and the positions of the dsb95 segments for different kinds of events.

### DSB sites are found between noncrossover trans hDNA tracts, which supports double SDSA and/or dHJ dissolution, and allows to infer strand orientation

Noncrossovers characterized by two hDNA tracts in a trans configuration on the same chromatid are expected to result either from double SDSA or from dHJ dissolution (Figure 1B-C and 2D). This implies that a single Spo11-DSB initiates them, and this DSB is necessarily 5’ of the converted strands of the hDNA tracts. Our observations support this hypothesis since we found that a DSB region overlaps the region between the hDNA tracts in 53 out of 57 cases (#222-278) (Figure 2D and S4), the remaining cases representing a fraction of events not significantly different from the disregarded fraction of DSBs (*p*-value = 0.53, exact binomial test). Consequently, we used such events to infer the orientation of the corresponding DNA strands, a robust and critical piece of information not available from mere sequence analysis (Figure S6).

### Both one-sided and two-sided noncrossovers support the SDSA pathway and reveal a significant fraction of events associated with template switching

After inference of strand orientation for 100 out of 185 SDSA-compatible events (#14-198, Figure S4), we found that 85 oriented events contain a DSB region 5’ of the converted strands of the hDNA tracts (Figure 2B) as expected from the canonical model. The 15 remaining events do not, which is statistically different from the expected 5% of events arising from the disregarded fraction of DSBs (*p* = 0.00014, exact binomial test). Unconverted SNPs between the presumed position of DSBs and an SDSA hDNA tract can result from template switching, with the invading end first engaging into the sister chromatid prior to engaging into the homolog (Figure 4A). Following this hypothesis, we estimate that about 10 out of the 15 events correspond to template switching, giving a probability of ~ 0.1 for a DNA end to switch template when initiating SDSA. This calculation is an underestimate since we consider that a causal DSB occurred at the expected location if this site overlaps a dsb95 region. However, the causal DSB could occur at a different position in the dsb95 region that could be compatible with template switching. In addition, template switching can take place in regions devoid of markers and be undetectable.

**Figure 4:**
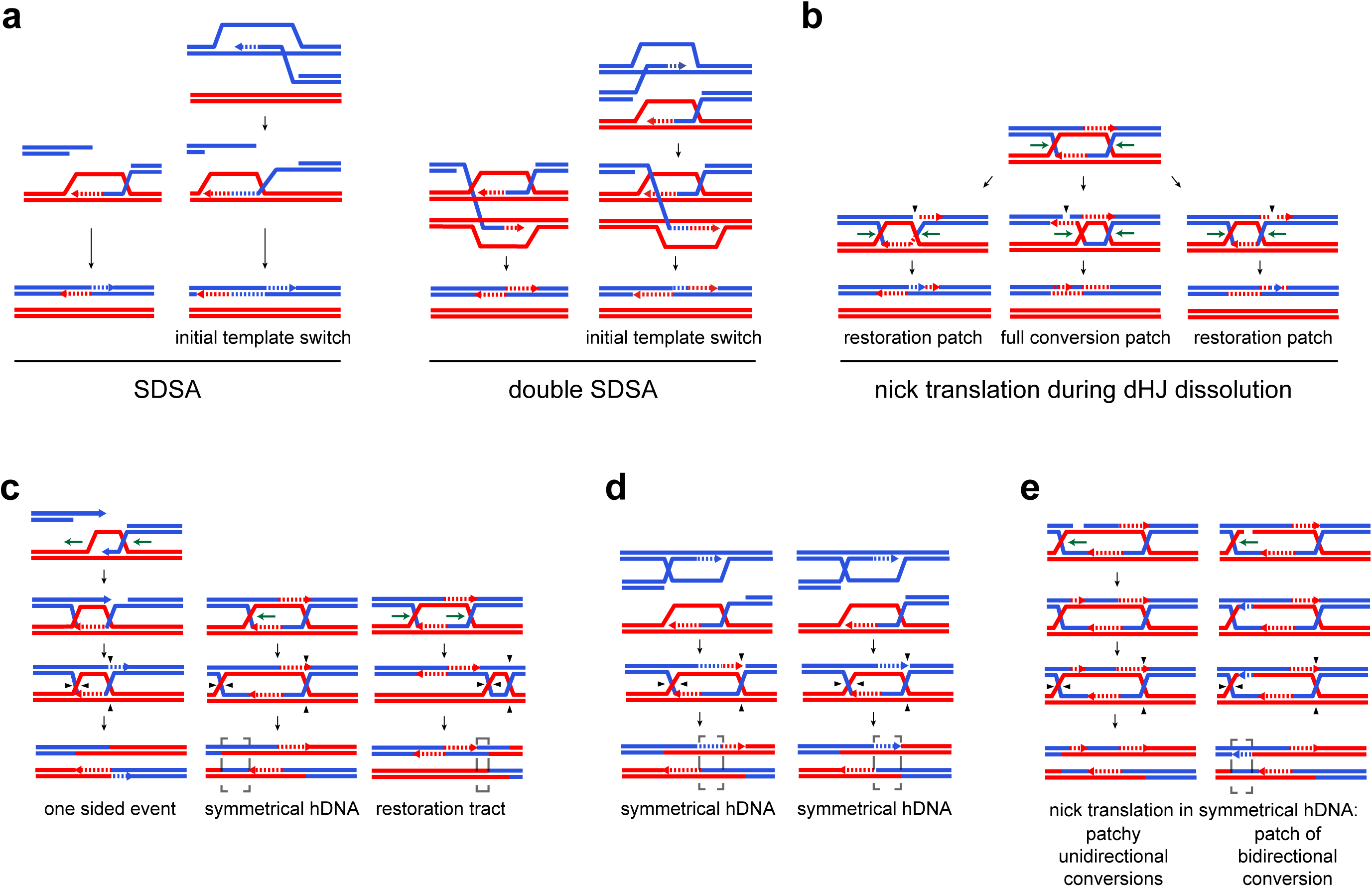
Variations from the canonical noncrossover and crossover formation pathways. **a**. The outcome of SDSA is a half-conversion tract located exactly 3’ of the invading end on the broken chromatid. In case of initial invasion of the sister chromatid prior to a switch to a non-sister chromatid, the half-conversion tract is separated from the 3’ invading end by a restoration tract. During single SDSA, this can result in the presence of unconverted SNPs between the DSB site and the halfconversion tract. During double SDSA, this automatically yields a restoration tract between two trans hDNA tracts. **b**. Nick translation during dHJ dissolution can also generate restoration tracts between trans hDNA tracts (left). However, depending on the location of the nick, nick translation can result in either restoration or full conversion, anywhere along the two trans hDNA tracts (middle and right). **c**. Branch migration and crossover resolution. Branch migration (green arrows) of crossover intermediates affects the strand transfer patterns of the recombination outcomes. Left. D-loop migration erases the hDNA tract composed of the invading 3’ end. Only one hDNA tract positioned on the left side of the initiating DSB is present in the final “one-sided” crossover. Middle. Branch migration of a single HJ away from the initiating DSB within a homoduplex DNA region generates symmetrical hDNA tracts (bracketed region). Right. Branch migration of the two HJ in the same direction can position the two hDNA tracts on the same chromatid while they were initially on the two recombining chromatids. Note that in case of noncrossover resolution, this generates trans hDNA tracts indistinguishable from those resulting from double SDSA or dHJ dissolution. In addition, in case of a long enough branch migration, a restoration tract can be formed between the hDNA tracts and the crossover site (bracketed region). Finally, symmetrical hDNA tracts corresponding to the region between the two HJs can also form. **d**. Template switching and crossover resolution. Invasion and DNA repair synthesis from a sister chromatid of one of the two 3’ ends prior to interacting with the displaced strand of the D-loop formed with the homologous chromatid by the other 3’ end generates a patch of symmetrical hDNA. This patch of symmetrical hDNA is flanked either by two hDNA tracts if there is DNA synthesis from the non-sister chromatid after template switch (left), or by a single hDNA tract in the absence of DNA repair synthesis from the non-sister chromatid (right). **e**. Nick translation within symmetrical hDNA tracts yields patchy unidirectional and bidirectional conversions.

Interestingly, among the 54 noncrossovers with trans hDNA tracts on the same chromatid showing additional restoration and full-conversion tracts (#279-332, Figure S4), 31 have a restoration patch exactly between the two hDNA tracts (#284-314, Figures 2E and S4). The over-representation of restoration over full-conversion patches and their biased location between the trans hDNA tracts support the hypothesis that these events result from the double SDSA pathway. Indeed, initial template switch between sister and non-sister chromatids automatically yields a restoration patch between the trans hDNA tracts (Figure 4A). In contrast, accumulation of nicks during dHJ dissolution followed by nick translation yields equal proportions of restoration and full-conversion patches within the hDNA tracts but not necessarily between the trans hDNA tracts (Figure 4B)^15^. Accordingly, comparing these 31 noncrossovers with a single restoration patch exactly between the two hDNA tracts with the 57 noncrossovers containing only two trans hDNA tracts (#222-278, Figure S4) allowed us to propose that the proportion of DNA tails performing template switching when initiating SDSA is likely higher than 0.2 (see Methods).

Overall, we bring unprecedented support to the SDSA model during meiosis. Interestingly, our template switching estimate is close to the estimate of template switching during break-induced replication in mitotic cells^41, 42^, a process sharing the same initial steps with SDSA. The frequent interaction with the sister is rather unexpected because of a strong homolog bias during meiotic recombination^43^. Initial invasion of the sister chromatid could be favored by the tight sister chromatid cohesion mediated by Rec8 and later released by Red1/Mek1^44^. Importantly, the heteroduplex rejection property of the mismatch repair machinery that could promote recombination with the sister chromatid is absent in our conditions ^45^. Therefore, template switching likely reflects an intrinsic property of the recombination mechanism that might be a consequence of the “collision release” process in which pol delta ejects from PCNA upon extending a DNA template by running into a downstream duplex^46^. Such template switching was proposed to promote the efficiency of the late steps of recombination^17^, and might also explain why most noncrossovers-associated gene conversions in mice are away from the expected DSB location^47, 48^.

### About one fifth of crossovers occur away from a mapped DSB region in events with simple patterns

The canonical DSB repair model predicts a strict connection between DSBs and hDNA tracts associated to crossovers (Figures 1C and 3A-H). Remarkably, for 46 (11%) out of the 412 crossovers involving two non-sister chromatids, the sequence located between the adjacent 4:4 segregating markers of the event (Figure S2A) does not overlap a dsb95 segment (Figures 3I and S5). This is statistically higher than the expected 5% (*p*-value = 6×10^‒7^, exact binomial test). This discrepancy between expected and observed DSB locations reaches one out of five cases for classes of crossovers with simple patterns where the expected DSB locations are more restricted, *i.e.* crossovers with no associated strand transfer, crossovers with a single full-conversion tract (#1-39, Figure S5), crossovers with a single hDNA tract with or without an additional full-conversion tract (#63-116 and #135-156, Figures S5 and 3C) (see Methods). Several possibilities could account for this observation, including template switching between sister and non-sister chromatids and HJ mobility.

### Extensive HJ migration during DSB repair

Out of 361 noncrossovers involving one or two non-sister chromatids, 29 (#333-361, Figure S4) have strand transfer on two homologous chromatids, which is expected from the resolution of a dHJ-containing intermediate (Figure 1C and 2H). Only two exhibit the hDNA pattern expected from canonical dHJ resolution (#333-334), while twenty contain symmetrical hDNA tracts. These tracts form either when a HJ migrates within its adjacent homoduplex DNA, or when a DNA end performs template switching engaging first into the sister chromatid prior to interacting with the homolog (Figure 4C and D). The high frequency of symmetrical hDNA tracts and the patchiness of the events that includes bidirectional conversion tracts (#353-361) support the following model: nicks form during HJ branch migration potentially through the abortive action of topoisomerases, and subsequent nick translation results in patches of unidirectional or bidirectional conversions (Figure 4B and E)^15^. HJ migration is also evidenced by noncrossovers #335-337 where inward migration of one HJ with respect to the initiating DSB likely splits the original hDNA tract in two pieces spread on the two recombining chromatids.

Likewise, many crossovers can be interpreted as exhibiting patterns characteristic of HJ migration, including the 102 bidirectional crossovers (#240-341, Figure 3D, F, J and S5), the 10 unidirectional crossovers with symmetrical hDNA tracts, and a few other events containing trans hDNA tracts on one chromatid (for example #199,200,204,223). In total, these crossovers represent at least 116 (28%) events out of the 412 crossovers involving two chromatids. Finally, the significant proportion of crossovers that do not exhibit a DSB region at the expected location can result from HJ mobility (Figure 3I).

In conclusion, HJs frequently migrate during crossovers and noncrossovers recombination. Although considered as a possibility by Szostak and colleagues^12^, this should now be included as a general feature of the DSB repair pathway and considered when studying recombination events lengths.

### Meiotic phenotypes of mutants affecting the ZMM pathway at different steps

The genetic control of the recombination patterns observed remains to be determined. We therefore studied the *msh5*Δ mutant thought to block the ZMM pathway early, the *mlh1*Δ, *mlh3*Δ, and *exo1*Δ mutants thought to block this pathway at the resolution step, the meiotic null p*CLB2*-*SGS1* mutant expected to affect several steps and pathways of recombination, and the *pms1*Δ and *mlh2*Δ mutants whose corresponding proteins form two additional complexes with Mlh1. Full viable *msh5*Δ octads could only be obtained in an *mlh2*Δ background^40, 49^.

Compared with the reference strain, we found a decrease in the number of crossovers in the *mlh1*Δ, *mlh3*Δ, *exo1*Δ, and *mlh2*Δ *msh5*Δ mutants consistent with previous reports^350-52^ (Figure 5A). This decrease seems more pronounced in the *mlh3*Δ, *exo1*Δ and *mlh2*Δ *msh5*Δ mutants that also show chromosomes segregating without crossovers (Table S1). The elevated number of noncrossovers in the *mlh2*Δ *msh5*Δ mutant is reminiscent of a pairing defect known to prevent negative feedback on DSB formation^53^. Finally, we found that the lengths of the DNA strand transfers associated to recombination events are increased to a similar extent in *mlhl*Δ and *mlh2*Δ mutants, further supporting that the Mlh1-Mlh2 heterodimer is required to limit hDNA tract extension^40^ (Figure 5B).

**Figure 5:**
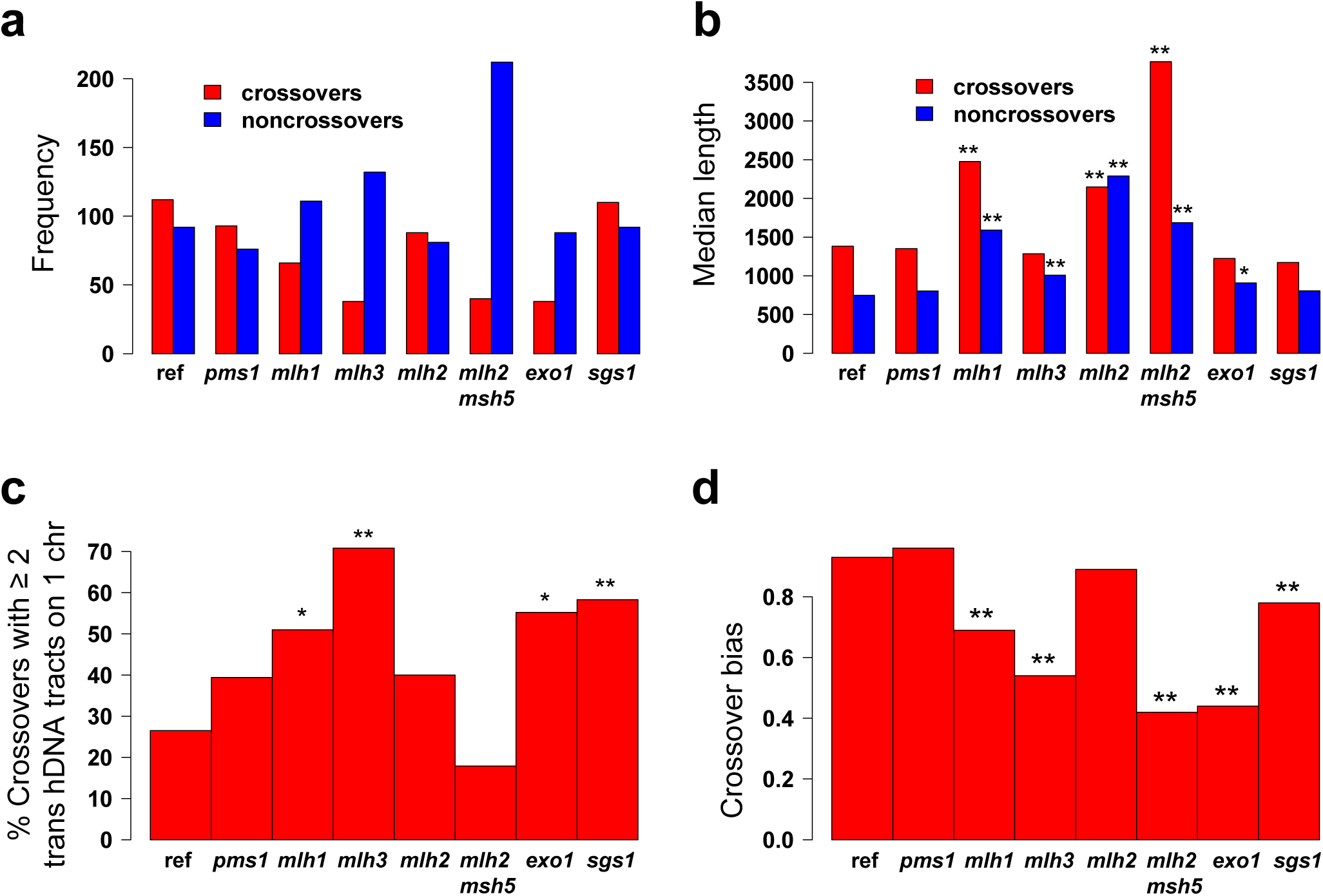
Meiotic characteristics of the strains used. **a**. Mean number of crossovers and noncrossovers per octad. **b**. Median length of the strand transfers associated with crossovers and noncrossovers. The length is defined as the distance between the midpoints of the intermarker intervals surrounding the recombination event. Stars indicate when the lengths are significantly different in the mutant and in the reference *msh2*Δ strain (one star, *p* < 0.05; two stars, *p* < 0.01, Wilcoxon test). For clarity reasons, the variability of the events lengths is not represented here but can be computed from Table S1. **c**. Two-sided crossovers subcategories. Percentage of two-sided crossovers with at least one pair of trans hDNA tracts on one chromatid. The stars indicate when the ratios are significantly different in the mutant and in the reference *msh2*Δ strain (one star, *p*-value < 0.05; two stars, *p*-value < 0.01, Fisher’s exact test). The numbers of events considered are 49, 33, 51, 24, 45, 28, 29 and 72 for the reference strain and the *pmsl*Δ, *mlhl*Δ, *mlh3*Δ, *mlh2*Δ, *mlh2*Δ *msh5*Δ, *exol*Δ and *sgsl* mutants, respectively. **d**. Crossover bias. Crossover bias corresponds to 1 minus the ratio between the noncrossovers derived from resolution and the total number of crossovers. Stars indicate when the crossover bias is significantly different from the reference *msh2*Δ strain (one star, *p* < 0.05; two stars, *p* < 0.01, Fisher’s exact test). The numbers of events considered are 470, 281, 165, 111, 189, 120, 117 and 257 for the reference strain and the *pmsl*Δ, *mlhl*Δ, *mlh3*Δ, *mlh2*Δ, *mlh2*Δ *msh5*Δ, *exol*Δ and *sgsl* mutants, respectively.

The most remarkable changes in noncrossovers concern p*CLB2*-*SGS1* (Figure 2C). The decrease in noncrossovers with trans hDNA tracts on one chromatid could reflect impaired joint molecule dissolution^54, 55, 56^ and would explain at least in part the increase in noncrossovers coming from dHJ resolution. The unchanged fraction of SDSA compatible events could result from functional redundancy with other helicases, such as Mph1 and Srs2^57^. It could also suggest that our strategy that relies on fully viable meiotic progeny is unable to reveal the potentially more abundant branched intermediates in the absence of Sgs1^17, 58^, or that the Sgs1 knock down is only partial in our condition^59^.

### The apparent processivity parameter of the DNA repair polymerase involved in the SDSA pathway is higher in the absence of Mlh1 or Mlh2

In the absence of Msh2-dependent mismatch repair, hDNA tracts resulting from SDSA correspond to tracts of newly synthesized DNA (Figure 1B). Since the distributions of the lengths of hDNA tracts associated to SDSA-compatible events are not statistically different for the reference and the *exo1*Δ strains (Figure 6), hDNA tract lengths are likely determined by the new strand synthesis only and not by the extent of Exo1-mediated resection^60, 61^. We therefore designed a strategy to extract the apparent processivity parameter of the DNA repair polymerase at work during SDSA (see Methods).

**Figure 6:**
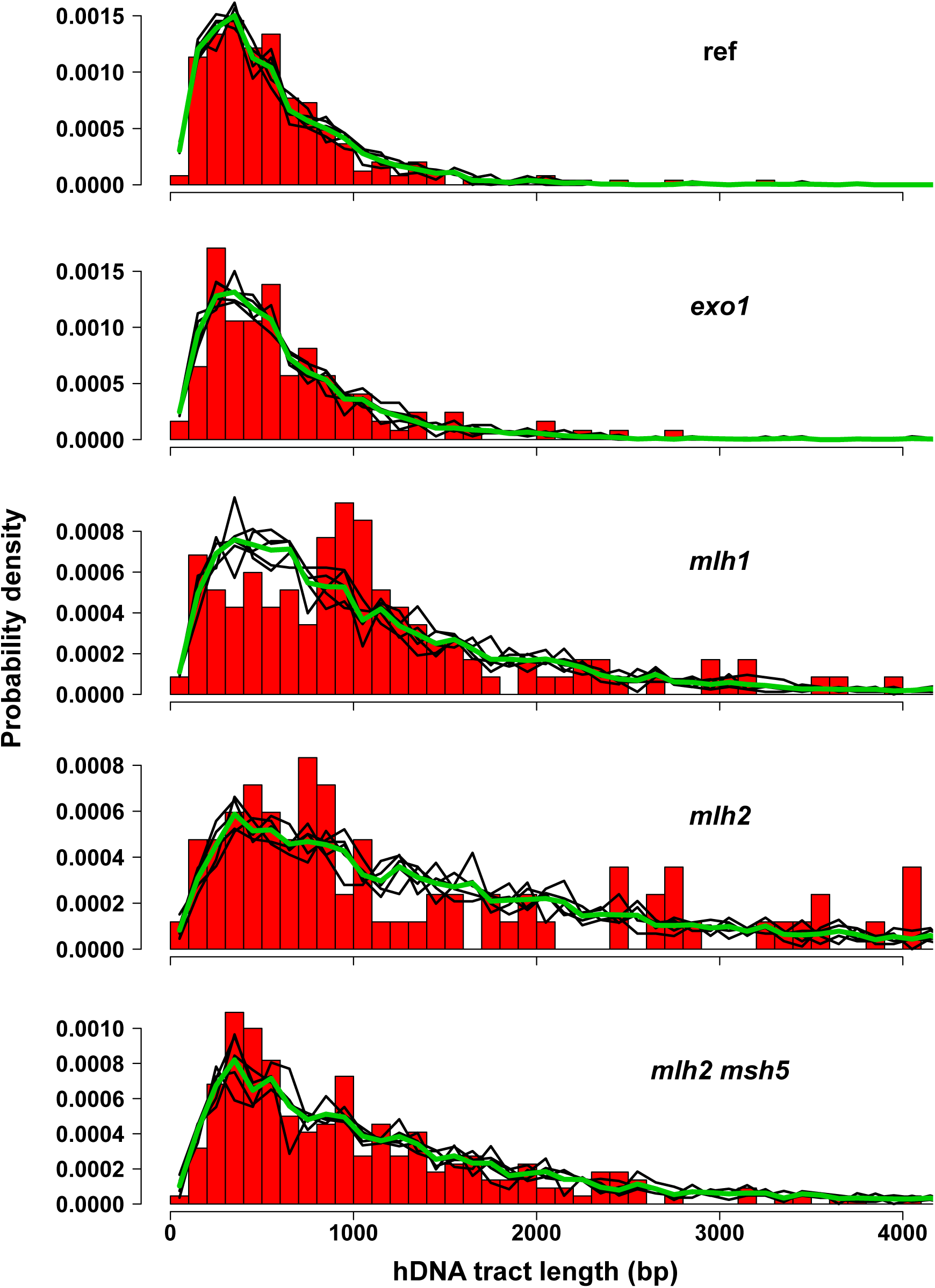
Distributions of hDNA tract lengths for the reference strain and several mutants. hDNA length corresponds here to *len*_*mid* values, defined as the distance between the midpoints of the intermarker intervals adjacent to the hDNA tract. The red histograms represent the distributions of the observed *len*_*mid* values divided into 100 bp bins. For each strain, the density functions of the simulation sets for the processivity value *p* giving the best fit to the observed distribution are shown as black, thin lines whereas the thick, green line represents the average of these density functions. The numbers of data values are 247, 123, 117, 84, and 220, for the reference strain and the *exol*Δ, *mlhl*Δ, *mlh2*Δ, and *mlh2*Δ *msh5*Δ mutants, respectively. Some extreme *len*_*mid* values are not represented on the graphs, namely 4510, 4841, and 5052 for the *mlhl*Δ mutant, 4220, 4504, 4750, 5048, and 7300 for the *mlh2*Δ mutant, and 4480, 4708, 5730, 6794, 7330, and 8884 for the *mlh2*Δ *msh5*Δ mutant.

Briefly, we resorted to simulations to test different models of polymerase activity. The choice of the distributions to model hDNA tracts lengths was guided by a basic model of processive enzyme according to which the polymerase has a probability *p* of moving to the next base and a probability *1*-*p* of falling off. For the reference strain, the best fit was obtained with *p* = 0.9972 (*p*-value = 0.52, chi-squared test) (Figure 6). The quality of the fit supported our model.

The distributions of hDNA tracts lengths for the *exo1*Δ, *mlh3*Δ, *pms1*Δ, and p*CLB2*-*SGS1* mutants were not different from that of the reference strain (*p*-value > 0.05, Fisher’s test). In contrast, the distributions of hDNA tracts lengths for the *mlhl*Δ, *mlh2*Δ, and *mlh2*Δ *msh5*Δ mutants were significantly different from the reference (*p*-value < 10^‒6^) with the best fits for *p* obtained with 0.9989 (*p*-value = 0.1), 0.9993 (*p*-value = 0.8), and 0.9990 (*p*-value = 0.7), respectively. The distributions of hDNA tracts lengths in the *mlh1*Δ, *mlh2*Δ, and *mlh2*Δ *msh5*Δ mutants can therefore be accounted for by the simple model of a polymerase whose processivity is markedly increased (0.9986-0.9994 compared to 0.9969-0.9977 for the reference strain).

Our data resemble the exponential relationship observed between the frequencies of gene conversion and the distance from the DSBs, interpreted as reflecting the processivity of a mismatch repair complex progressing from the DSB site with a probability *p* of moving on to the next nucleotide. Data from the *ARG4* locus led to a broad estimate of *p* between 0.9975 and 0.999 (for a review see^62^). However, comparison with our data is difficult since in an Msh2-dependent mismatch repair-proficient context, conversion tracts are the end results of a series of processes including DNA synthesis and mismatch repair of heteroduplexes that could occur at several steps and lead either to conversion or restoration. Our *msh2*Δ background constitutes a simpler system that allowed for the first time, to our knowledge, to determine the apparent *in vivo* processivity parameter of a DNA repair polymerase.

### Role of Mlh1-3 in the biased resolution of crossover-intermediates

The two crossover resolution configurations can be distinguished depending on whether the D-loop invading strand is cleaved or not (Figure 1C and S7, see Methods). From the parsimonious analysis of four *msh2*Δ reference octads, 51 onesided and 27 two-sided crossovers showed a resolution pattern #1, and four onesided and zero two-sided crossovers showed a resolution pattern #2. The ratio of crossovers with pattern #2 is significantly different from 0.5 in both cases (*p*-value equal to 2.0×10^‒11^ and 1.5×10^‒8^, respectively, exact binomial test), revealing a strong bias toward resolution pattern #1 (Table 1). While such a bias was clearly established during mitotic recombination in mammalian and yeast cells^63–65^, it was also detected during yeast and fly meiosis at a few hot spots but with no definitive conclusion^33, 34, 37, 66^. The nature of the signal associated with the newly synthesized DNA is unknown, but an attractive hypothesis is that the nicks left after DNA repair synthesis are not ligated prior resolution. Resolution of dHJs as crossovers would therefore require only two additional nicks instead of four. This would also tend to bias resolution toward crossovers versus noncrossovers, since noncrossover resolution would require three nicks. Existence of nicked HJs was suggested from electron microscopic observation of chromosomal DNA from pachytene yeast cells^23, 67^, but is not supported by the work of Schwacha and Kleckner that observed a majority of fully ligated joint molecules at the *HIS4*-*LEU2* hotspot^43^. Remarkably, the double-stranded DNA nicking activity of Mlh1-Mlh3 would allow resolution of nicked HJs much better than of fully ligated HJs^19–21^.

**Table 1.** Numbers of one-sided and two-sided crossovers resolved with patterns #1 and #2. pvalue 1_0.5 and pvalue 2_0.5: *p*-values corresponding to the exact binomial test that the ratio of crossovers with pattern #2 is not statistically different from 0.5, for one-sided crossovers and for two-sided crossovers, respectively. pvalue 1_ref and pvalue 2_ref: *p*-values corresponding to Fisher’s test that the ratio of crossovers with pattern #2 in the reference and in the mutant strains are not statistically different, for one-sided crossovers and for two-sided crossovers, respectively.

|  | one-sided | one-sided | two-sided | two-sided | pvalue | pvalue | pvalue | pvalue |
| --- | --- | --- | --- | --- | --- | --- | --- | --- |
| genotype | pattern 1 | pattern 2 | pattern 1 | pattern 2 | 1_0.5 | 2_0.5 | 1_ref | 2_ref |
| ref | 51 | 4 | 27 | 0 | $2.0 \times 10^{-11}$ | $1.5 \times 10^{-8}$ | 1.0 | 1.0 |
| *pms1* | 26 | 2 | 19 | 0 | $3.0 \times 10^{-6}$ | $3.8 \times 10^{-6}$ | 1.0 | 1.0 |
| *mlh1* | 13 | 7 | 30 | 1 | 0.26 | $3.0 \times 10^{-8}$ | 0.0061 | 1.0 |
| *mlh3* | 3 | 4 | 7 | 0 | 1.0 | 0.016 | 0.0037 | 1.0 |
| *mlh2* | 20 | 3 | 29 | 0 | 0.00049 | $3.7 \times 10^{-9}$ | 0.41 | 1.0 |
| *mlh2 msh5* | 6 | 2 | 19 | 1 | 0.29 | $4.0 \times 10^{-5}$ | 0.16 | 0.43 |
| *exo1* | 6 | 0 | 6 | 0 | 0.031 | 0.031 | 1.0 | 1.0 |
| *sgs1* | 15 | 2 | 20 | 1 | 0.0023 | $2.1 \times 10^{-5}$ | 0.62 | 0.44 |

We found that in all the mutants tested the proportion of two-sided crossovers with pattern #2 over the total number of two-sided crossovers was not different from that of the *msh2*Δ reference strain and that they all exhibit a clear bias in favor of pattern #1 (Table 1). In contrast, we observed that the proportion of one-sided crossovers with pattern #2 over the total number of one-sided crossovers was significantly different from that of the reference strain for the *mlh3*Δ and *mlh1*Δ mutants. This suggests that Mlh1-3 specifically promotes pattern #1 dHJ resolution of one-sided crossover intermediates. Consequently, the alternative resolution pathway mainly relying on Mus81 recognizes the imprints associated to the newly synthesized DNA only for two-sided crossover intermediates. If these imprints are nicks, they should be recognized and lead to crossover after Mus81 resolution^68, 69^). The difference between one-sided and two-sided intermediates suggests that either such nicks are lost in the absence of Mlh1-3 specifically for one-sided crossover intermediates, or that the imprints are not nicks.

### Mlh1-3, Mlh1-2, Exo1 and Sgs1 promote one-sided crossovers with bidirectional conversions and prevent accumulation of symmetrical hDNAs

In the *mlhl*Δ, *mlh3*Δ, *exo1*Δ, *mlh2*Δ, p*CLB2*-*SGS1* and *mlh2*Δ *msh5*Δ mutants, we found a global trend for crossover-associated strand transfers, that includes a decrease in one-sided bidirectional gene conversions compared to the *msh2*Δ reference strain, an increased frequency of two-sided crossovers and an increase in symmetrical hDNA tracts (Figure 3J and S8).

Bidirectional gene conversions have always been considered as a result of multiple initiating Spo11-DSBs in trans, normally repressed by Tel1 and Mec1^70^. Under this scenario, our observation would mean that Mlh1, Mlh2, Mlh3, Exo1 and Sgs1 promote localized Spo11-DSBs in trans, although these proteins have no known effect on Spo11 activity. Remarkably, bidirectional events mostly involve only two chromatids, while a significant fraction of events involving at least three chromatids would be expected if they resulted from multiple independent initiating Spo11-DSBs. This leads to the alternative hypothesis postulated above where bidirectional events would reflect Spo11-independent DNA breaks occurring during the processing of recombination intermediates (Figure 4E). Mlh1-3, Mlh1-2, Exo1 and Sgs1 would promote such processing, where the nuclease activity of Mlh1-3 could be a source of nicks and even DSBs directly generating bidirectional conversions^19–21^. The decrease in one-sided bidirectional conversions paralleled with the increase in two-sided crossovers and crossovers with symmetrical hDNA tracts could suggest a connection between these patterns controlled by Mlh1-3, Mlh1-2, Exo1 and Sgs1. Importantly, this result combined with other recent results showing that Mlh1-2 limits the extent of meiotic hDNA tracts^40^ and that Mlh2 was lost concomitantly with Zip2, 3, 4, Spo16 and Msh4-5 in *Lachancea* yeasts ^71^, reinforces the tight connection between Mlh1-2 and the ZMM pathway.

Regarding two-sided crossovers, the *mlh1*Δ, *mlh3*Δ, *exo1*Δ and p*CLB2*-*SGS1* mutants have more frequently trans hDNA tracts located on one chromatid rather than on two recombining chromatids (Figure 3H, 5C). This pattern is reminiscent of the resolution of a dHJ-containing intermediate <u>*after*</u> the migration of one HJ toward the other HJ beyond the DSB site (Figure 4C) ^72^. This result suggests that Mlh1, Mlh3, Exo1 and Sgs1 limit inward HJ branch migration. Besides, the increase in symmetrical hDNA tracts in their absence suggests that these four proteins also limit outward HJ branch migration (Figure 5C). Accordingly, Sgs1 seems to prevents branch migration under some circumstances, while this helicase is so far known to undo joint molecules^54, 55, 56^. The polymer structure of Mlh1-3 might be important to limit this Sgs1-independent HJ migration^19^. Moreover, these results support a possible unanticipated role of Mlh1-3 before the resolution step^51^.

### Crossover resolution bias of dHJs

Considering that noncrossovers involving two non-sister chromatids derive from dHJ resolution, we could directly measure the crossover resolution bias of dHJs previously identified^28–30^. We expressed it as 1 minus the ratio between noncrossovers resulting from dHJ resolution and the total number of crossovers. Compared to the *msh2*Δ reference, the bias is not affected in the *pms1*Δ nor *mlh2*Δ mutants, but is significantly decreased in the p*CLB2*-*SGS1*, *mlh1*Δ, *mlh3*Δ, *exo1*Δ and *mlh2*Δ *msh5*Δ mutants (Figure 5D). It is tempting to speculate that the crossover bias could result from the dHJ resolution bias. However, the dHJ resolution bias is always maintained for the two-sided crossover intermediates while the *exol*Δ, *mlhl*Δ, *mlh3*Δ and *mlh2*Δ *msh5*Δ mutants show a strong crossover bias defect. This shows that the crossover bias and the dHJ resolution bias are independent, the crossover bias being established prior to resolution.

## DISCUSSION

The comprehensive analysis of meiotic hDNA from cells lacking the Msh2 mismatch repair factor clarified our understanding of the mechanistic of meiotic recombination. It revealed a remarkable and so far under appreciated dynamics of the meiotic DNA transactions with frequent template switching and branch migration. It confirmed the core of the DSB repair pathway, but identified variations from its canonical version. The comparative analysis of different mutants notably revealed that Mlh1-3 promotes both the asymmetric maturation and the biased resolution of crossover intermediates, properties in line with its recent functional characterization^19–21, 50, 51^. Given the conservation of the proteins and pathways, we expect our findings apply to many eukaryotes including mammals.

## AUTHOR CONTRIBUTIONS

Conceptualization, MC.MK. and B.L.; Methodology, MM.K., MC.MK. and B.L.; Software, MC.MK.; Formal Analysis, MC.MK.; Investigation, MM.K., J.S., X.Z., MC.MK. and B.L; Data Curation, MC.MK. and B.L.; Writing – Original Draft, MC.MK. and B.L.; Supervision, MC.MK. and B.L.; Funding Acquisition, B.L.

## ACKNOWLEGMENTS

MCMK would like to thank Bruno Toupance (MNHN, Paris) for helpful discussions on probability. We thank Scott Keeney for providing Spo11-oligo data, Olivier Espeli, Romain Koszul and Mauro Modesti for discussions and Emmanuelle Martini and Scott Keeney for critical reading of the manuscript. BL lab is funded by the ANR (Agence nationale de la recherche) ANR-13-BSV6-0012-01 and ANR-16-CE12-0028-01 grants. MMK is a recipient of a post doctoral fellowship from La Ligue Contre le Cancer. JS was a recipient of a doctoral fellowship from La Ligue Contre le Cancer. XZ was supported by National Institutes of Health grant R01 GM058673 (to S.Keeney).

## MATERIALS AND METHODS

### Strains and media

All yeast strains used in this study are derivatives of S288C^73^ and SK1^74^. Strain genotypes are listed in Table S2. Gene disruptions were performed by PCR-mediated gene replacement^75^ followed by PCR analysis for discriminating correct and incorrect gene targeting. S288C × SK1 crosses were made on YPD plates and diploids were subcloned onto selective plates before transfer on 1% potassium acetate sporulation medium and subsequent dissection. The hybrids made for octad sequencing are BLY-HY1 to 8 (Table S2). Note that the octad datasets for *msh2*Δ (BLY-HY1) and *msh2*Δ *mlh2*Δ (BLY-HY5) are the same as in^40^.

### Spo11-oligo mapping

A previously described, fully functional FLAG-tagged version of Spo11 was used^53^, ^76^. The starting S288C strain was strain YTT0559 from the T. Tsukiyama lab, Fred Hutchinson Cancer Research Center and was generously provided by I. Whitehouse, Memorial Sloan Kettering Cancer Center. The endogenous *SPO11* locus in this strain was tagged by integration of a *6His*-*3FLAG*-*loxP*-*kanMX*-*loxP* construct amplified from an SK1 strain provided by Kunihiro Ohta, Univ. of Tokyo^76^, creating SKY4300. Correct tagging was verified by PCR and Southern blot. The SK1 parent (SKY4301) was derived from standard SK1 strains in the S. Keeney laboratory (Memorial Sloan Kettering Cancer Center) and has a series of single-base changes introduced within and near the DSB hotspots in the *PDR3*, *RIM15*, *CCT6*, and *IMD3* promoters; these base changes do not detectably alter the DSB distribution as assessed by Spo11-oligo mapping (X. Zhu, S. Globus, and S. Keeney, unpublished observations). SKY4300 and SKY4301 were mated to generate the F1 hybrid diploid strain (SKY4302) used for Spo11-oligo mapping.

Spo11-oligo mapping was performed as previously described^77^, ^78^ except that meiotic cultures from SKY4302 were harvested 6 hr after transfer to sporulation medium. Briefly, Spo11 oligos were purified by immunoprecipitation with anti-FLAG antibodies, then sequencing adapters were added and the oligos were amplified by PCR and sequenced. Sequencing was performed using Illumina HiSeq in the Memorial Sloan Kettering Cancer Center Integrated Genomics Operation core facility. *In silico* clipping of sequencing adapters and mapping of reads to the sacCer2 genome assembly was performed using a previously described custom pipeline^53^. After mapping, the reads were separated into unique and multiple mapping sets, but only uniquely mapping reads were used in this study (multiple mapping reads constituted a small minority of the total). In Figures 2–3 and S6, the absolute number of uniquely mapping Spo11-oligos is shown for each position.

### DNA extraction and octad sequencing

To determine the meiotic strand transfers that occurred in the absence of Msh2, we systematically generated and genotyped octads from S288C × SK1 hybrids. As described in Figure S1, octads were obtained by inducing sporulation of such hybrids and separating the mother from the daughter cells after the first mitotic division of the four viable spores. Genomic DNA was purified from overnight saturated YPD culture using a Qiagen genomic-tip 100/G. Sequencing was performed at BGI using Illumina HiSeq instruments. We analyzed four octads from the *msh2*Δ hybrid, three from the *msh2*Δ *pmsl*Δ hybrid and two from all the other hybrids. For each genetic background, all the recombination events from all the octads sequenced were pooled for analysis.

### Genotyping and recombination event calling

Spores were sequenced to an average sequencing depth of 50-60x. The reads were aligned on the reference sequences of the parental strains S288C and SK1 using the BWA software^79^. We used the sequences from the *Saccharomyces* Genome Database (SGD) web site (http://www.yeastgenome.org/) for S288C and the sequence made available by S. Keeney (http://cbio.mskcc.org/public/SK1MvO/) for SK1. Only the reads matching perfectly the reference sequences were taken into account. S288C-SK1 genome alignment was performed using the LAGAN software^80^. Only the aligned blocks longer than 2 kb were taken into account and within these blocks only single nucleotide polymorphisms (SNPs) were considered. The parental strains used in this study were also sequenced and all positions whose base identity was not confirmed were eliminated. Finally, SNPs located in repeated regions, long terminal repeats (LTRs), retrotransposons and telomeres were discarded, which yielded a final list of 74,911 SNPs.

Genotyping was performed using stringent criteria: for a given SNP position, if reads matched only one parental sequence, at least 15 reads were required, but if reads aligned to both parental sequences, less than 5 reads matching one parental sequence and at least 40 reads matching the other parental sequence were required for base identity to be confirmed. Chromosomes were then segmented into stretches of SNPs with identical genotypic patterns. These patterns are coded as *x:y* with *x* representing the number of S288C alleles and *y* = *8*-*x* representing the number of SK1 alleles. Accordingly, a 4:4 segment corresponds to a Mendelian segregation profile, 5:3 and 3:5 segments to half-conversion tracts, 6:2 and 2:6 segments to full-conversion tracts, etc.

We defined a recombination event as a set of adjacent segments located between two 4:4 segments longer than 1.5 kb (the length used here is the distance between the midpoints of the intermarker intervals adjacent to the segment). It has to be noted that one recombination event can contain zero, one or two crossovers. Figure 5A gives the total number of crossovers present in the events (counting two crossovers for the few events concerned). In the rest of the article “crossover” refers to “a recombination event including at least one crossover”. The events were classified as a function of the number of chromatids involved: one chromatid, two chromatids, either sister or non-sister, and more than two chromatids. Only events affecting one chromatid or two non-sister chromatids are represented in Figures 2, 3, S4 and S5.

Overall, we tried to come up with the most parsimonious explanation for the strand transfer tracts observed at recombination events. The complexity of strand transfer tracts includes restoration patches, full-conversion patches and bidirectional conversions. A simple scenario to explain restoration and full-conversion patches consists of invoking an Msh2-independent short-patch mismatch repair pathway^81^ but there is no evidence in *S. cerevisiae* that such a pathway exists independently of recombination. Accordingly, throughout the paper and when possible, we tried to propose more parsimonious recombination-linked mechanisms to explain these patches, whose occurrence would be independent from mismatch repair processes. For instance, restoration tracts are compatible with template switch between the sister and the non-sister chromatids (Figure 4A) and full conversions can be the scars of gap repair (Figure S9). We also propose that restorations, full conversions and bidirectional conversions could result from a nick translation process, initiated at nicks that would form during the processing of recombination intermediates ^15^ (Figure 4B and 4E). These nicks could result from aborted actions of SSNs or topoisomerases such as Top3 during HJ branch migration, but also from Mlh1-3 that can even generate DSBs^19^. Finally, some of the full conversions observed might also derive from the mismatch repair-independent action of the proof-reading activity of polymerase delta^82^.

### Estimating the frequency of initial template switching in the SDSA pathway

The frequency of initial template switching between sister and non-sister chromatids during SDSA can be estimated by comparing the 31 noncrossovers with a single restoration patch exactly between the two hDNA tracts (that likely derive from double SDSA with at least one of the DNA ends performing template switching) with the 57 noncrossovers containing only two trans hDNA tracts (#222-278, Figure S4). If these 57 noncrossovers all derive from dHJ dissolution, and none from double SDSA, we get a probability of 1 for the DNA ends to perform template switching when initiating SDSA. Conversely, if these 57 noncrossovers all derive from double SDSA, we can calculate the probability *p* of initial template switching by one of the DNA ends when initiating SDSA as follows. The proportion of SDSA events without template switching, 57/(57 + 31), corresponds to the probability that none of the two DNA ends will perform template switching, that is *(1*-*p)(1*-*p).* From this, we can extract the value of *p* = 0.20. Because of the probable existence of undetected central restoration patches, this calculation yields an underestimate of *p*, so we propose that the proportion of DNA tails performing template switching when initiating SDSA is likely higher than 0.2.

### Comparing expected and observed DSB locations for crossovers with simple patterns

We consider classes of crossovers with simple patterns, for which the DSB locations that are expected under the standard recombination models are restricted to one segment or even one intermarker interval. There are no overlaps between DSB regions (dsb95 segments) and event sectors for 16 (23%) out of 71 crossovers with no associated strand transfer (Figure 3A and I), nor for 7 (18%) out of 39 crossovers with a single full-conversion tract (#1-39, Figure S5 and 3B) that are compatible with gap repair. In both cases, the ratios are statistically different from the expected 5% (*p*-values equal to 4×10 and 3×10, respectively). For some events, determining chromatid orientation allows to define more precisely the expected location of DSB. We were able to infer chromatid orientation for 55 events among 54 crossovers with a single hDNA tract (#63-116, Figures S5 and 3C) and 22 crossovers consisting of a single full-conversion tract and a single hDNA tract (#135-156, Figure S5). No DSB region was present at the expected location in 13 (24%) cases out of these 55 oriented crossovers (*p*-value = 2×10^‒6^).

### Estimating the apparent processivity parameter of the DNA repair polymerase involved in the SDSA pathway

Several options are available for determining the mechanistic parameters of the DNA repair polymerase involved in the SDSA pathway. A first option is to try to extract from the markers positions of each event an estimate of the actual length of the hDNA tract that includes the newly-synthesized DNA strand. The parameter *len*_*mid*, corresponding to the distance between the midpoints of the intermarker intervals adjacent to the hDNA tract, is often used as such an estimate. However this estimate exhibits several short-comings. First, it tends to be artificially inflated for short hDNA tracts. Second, the value of *len*_*mid* crucially depends on the local density of markers. Since the markers density widely fluctuates along the genome and could be biased relative to the positions of the Spo11-DSB hotspots, it is difficult to find a model allowing to extract from *len*_*mid* an adjusted estimate of the actual hDNA lengths.

We therefore resorted to simulations for testing models of the polymerase activity by comparing the distributions of observed and simulated *len*_*mid* values. We calculated the *len*_*mid* values of two sets of noncrossovers enriched in SDSA events: the noncrossovers comprising a single hDNA tract that we consider most likely derived from the single SDSA pathway (#14-198, Figures 1B, 2B and S4 for the reference strain) and the noncrossovers with a restoration patch exactly between two trans hDNA tracts that we consider as probably originating from the double SDSA pathway (#284-314, Figures 4A, 2E and S4 for the reference strain). In all cases, for a given strain, the distributions of the *len*_*mid* values of the two sets of noncrossovers were not statistically different and all *len*_*mid* values were therefore pooled together.

We considered the simplest model of a processive enzyme with a probability *p* of moving to the next base and a probability *1*-*p* of falling off. Sets of 1,000 DSB positions were sampled across the genome, weighting each position by its frequency of Spo11-DSBs, as determined by the counts of immunoprecipitated Spo11-FLAG oligos. Random values of hDNA lengths were generated from exponential distributions of parameter *1*-*p.* Different ranges of values were tested for *p*, depending on the strains. For each value of *p*, independent sets of 1,000 simulated *len*_*mid* values were generated. From the genomic boundaries of the simulated hDNAs (associating a sampled DSB position and a random value of hDNA length) and the positions of the local markers, we determined *len*_*mid* values. The distribution of the observed *len*_*mid* values was then compared to the average density function of five independent sets of simulated *len*_*mid* values, using Pearson’ chi-squared test. Considering only the values of *p* for which the *p*-value for Pearson’s goodness-of-fit test was higher than 0.1, we could estimate the processivity of the repair polymerase between 0.9969 to 0.9977 for the reference strain, between 0.9971 and 0.9982 for the *exol*Δ mutant, between 0.9990 and 0.9994 for the *mlh2*Δ mutant and between 0.9986 and 0.9991 for the *mlh2*Δ *msh5*Δ mutant.

Finally we found that for the reference strain the distribution of the lengths of the 114 hDNA tracts associated to the 57 noncrossovers composed of two trans hDNA tracts on the same chromatid (#222-278, Figures 2D and S4) was not statistically different from that of the single SDSA-compatible events (*p*-value = 0.62, Fisher’s test). This suggests either that the noncrossovers with two trans hDNAs on the same chromatid derives mostly from double SDSA or, if these noncrossovers derive from dHJ dissolution, that the processivity of the polymerase operating in this second pathway is similar to that of the polymerase operating during SDSA.

### Chromatid strand orientation

The reconstruction of the sequences of the two DNA strands of each chromatid in a given tetrad by sequencing the eight genomes of the corresponding octad does not allow to infer the strand orientation of the chromatids from the tetrad (Figure S1). To extract chromatid strand orientation, we performed several steps using recombination events with patterns of increasing complexity. In a first step, we used noncrossovers involving only one chromatid and including trans hDNA tracts, either adjacent or separated by a single restoration or full-conversion tract. Chromatid orientation could be automatically inferred from these events since the initiating DSB is expected to be located 5′ of the converted strand of the hDNA tracts (Figure 1B). In a second step, chromatid orientation was manually derived from the analysis of crossovers and noncrossovers involving two non-sister chromatids and including trans hDNA tracts, either adjacent or separated by a single restoration or full-conversion tract, in addition to crossovers containing only one half-conversion tract and one full-conversion tract (that correspond presumably to gap repair (Figure S7)). At that stage strand orientation determination benefited from the information gathered at the first step and from the Spo11-oligo-based DSB map. Finally, crossovers and noncrossovers containing symmetrical hDNA tracts were used to automatically complete chromatid orientation since in these cases the strand orientation of one chromatid allows to infer the strand orientation of the other chromatid involved. For each octad, 4 × 16 chromatids had to be oriented. In many cases, chromatid orientation could be deduced from different events and we systematically checked the consistency of these inferences (Figure S6). For example, for the first octad of the reference strain, the orientation of 29 chromatids was determined more than once, and all multiple orientations were consistent. In other cases, events involving one of the few chromatids with inconsistent orientations were disregarded.

Chromatid orientation was used (i) for deducing the expected location of DSB regions for one-sided crossovers and noncrossovers, *i.e.* the 5′ end of the converted strand of the hDNA tract, and (ii) for determining the resolution pattern of crossovers.

### Determining the resolution pattern of crossovers

Crossover resolution of a dHJ from the canonical DSB repair pathway involves two nicks, and two nick combinations are possible (Figure 1C). The two resolution configurations can be distinguished depending on whether the D-loop invading strand is cleaved or not. When the D-loop invading strand is not cleaved, the two resulting hDNA tracts are in continuity with the two parental strands of the broken chromatid and consist of 3’ overhangs exclusively (pattern #1, Figure 1C). However, when the D-loop invading strand is cleaved, the two resulting hDNA tracts are separated from the two parental strands of the broken chromatid by a full-conversion tract (pattern #2, Figure 1C). Overall, this pattern #2 resolution is almost never observed (Table 1).

Importantly, when only one hDNA tract is detected, as frequently observed, the hDNA tract is automatically in continuity with the parental strands of the broken chromatid. In this latter case, resolution pattern #1 yields a 3’ overhang while pattern #2 yields a 5’ overhang, which allows discrimination between the two resolution patterns (Figure S7A and B). Further complexity can arise upon HJ migration. In addition to a 3’ overhang, pattern #1 can yield a 5′ overhang, while pattern #2 can yield a 3’ overhang in addition to a 5’ overhang (Figure S7C). In such a situation, the initiating DSB is on one side of the two hDNA tracts while it is between the hDNA tracts in the canonical DSB repair pathway. These patterns do not allow discrimination between resolution pattern #1 and #2.

Overall, resolution pattern #2 for one-sided events is inferred from the presence of a hDNA tract that is a 5’ overhang. However, resolution pattern #1 could also formally yield a 5’ overhang after HJ branch migration and when one hDNA tract is not detected as in Figure S7C. We disregarded this scenario because it is less parsimonious than the simple absence of one hDNA tract (Figure S7A), but we cannot formally exclude it.

### Statistical analyses

All statistical analyses were performed with the R environment (http://www.R-project.org/). Fisher’s test, Wilcoxon’s test and the exact binomial test, corresponding respectively to the R functions fisher.test(), wilcox.test(), and binom.test(), were always used with the two-sided option. Wilcoxon’s test is applied with a continuity correction.

### Data availability

Octad sequences are publicly available at the NCBI Sequence Read Archive (accession number SRP111430).

Sequence reads and compiled Spo11-oligo maps are publicly available at the Gene Expression Omnibus (GEO) repository (accession number GSE101339).

## REFERENCES

1. Keeney, S., Giroux, C. N. & Kleckner, N. Meiosis-specific DNA double-strand breaks are catalyzed by Spo11, a member of a widely conserved protein family. Cell 88, 375–384 (1997).

2. Sun, H., Treco, D., Schultes, N. P. & Szostak, J. W. Double-strand breaks at an initiation site for meiotic gene conversion. Nature 338, 87–90 (1989).

3. Zakharyevich, K. et al. Temporally and biochemically distinct activities of Exo1 during meiosis: double-strand break resection and resolution of double Holliday junctions. Mol. Cell 40, 1001–1015 (2010).

4. Sun, H., Treco, D. & Szostak, J. W. Extensive 3′-overhanging, single-stranded DNA associated with the meiosis-specific double-strand breaks at the ARG4 recombination initiation site. Cell 64, 1155–1161 (1991).

5. Mimitou, E. P., Yamada, S. & Keeney, S. A global view of meiotic doublestrand break end resection. Science 355, 40–45 (2017).

6. Garcia, V., Phelps, S. E. L., Gray, S. & Neale, M. J. Bidirectional resection of DNA double-strand breaks by Mre11 and Exo1. Nature 1–6 (2011). doi:10.1038/nature10515

7. Neale, M. J., Pan, J. & Keeney, S. Endonucleolytic processing of covalent protein-linked DNA double-strand breaks. Nature 436, 1053–1057 (2005).

8. Nassif, N., Penney, J., Pal, S., Engels, W. R. & Gloor, G. B. Efficient copying of nonhomologous sequences from ectopic sites via P-element-induced gap repair. Mol. Cell. Biol. 14, 1613–1625 (1994).

9. Ferguson, D. O. & Holloman, W. K. Recombinational repair of gaps in DNA is asymmetric in Ustilago maydis and can be explained by a migrating D-loop model. Proc. Natl. Acad. Sci. U.S.A. 93, 5419–5424 (1996).

10. Pâques, F. & Haber, J. E. Multiple pathways of recombination induced by double-strand breaks in Saccharomyces cerevisiae. Microbiol. Mol. Biol. Rev. 63, 349–404 (1999).

11. Resnick, M. A. The repair of double-strand breaks in DNA; a model involving recombination. J. Theor. Biol. 59, 97–106 (1976).

12. Szostak, J. W., Orr-Weaver, T. L., Rothstein, R. J. & Stahl, F. W. The doublestrand-break repair model for recombination. Cell 33, 25–35 (1983).

13. Gilbertson, L. A. & Stahl, F. W. A test of the double-strand break repair model for meiotic recombination in Saccharomyces cerevisiae. Genetics 144, 27–41 (1996).

14. Porter, S. E., White, M. A. & Petes, T. D. Genetic evidence that the meiotic recombination hotspot at the HIS4 locus of Saccharomyces cerevisiae does not represent a site for a symmetrically processed double-strand break. Genetics 134, 5–19 (1993).

15. Martini, E. et al. Genome-wide analysis of heteroduplex DNA in mismatch repair-deficient yeast cells reveals novel properties of meiotic recombination pathways. PLoS Genet. 7, e1002305 (2011).

16. Lynn, A., Soucek, R. & Börner, G. V. ZMM proteins during meiosis: crossover artists at work. Chromosome Res. 15, 591–605 (2007).

17. Oh, S. D. et al. BLM Ortholog, Sgs1, Prevents Aberrant Crossing-over by Suppressing Formation of Multichromatid Joint Molecules. Cell 130, 259–272 (2007).

18. Zakharyevich, K., Tang, S., Ma, Y. & Hunter, N. Delineation of joint molecule resolution pathways in meiosis identifies a crossover-specific resolvase. Cell 149, 334–347 (2012).

19. Manhart, C. M. et al. The mismatch repair and meiotic recombination endonuclease Mlh1-Mlh3 is activated by polymer formation and can cleave DNA substrates in trans. PLoS Biol. 15, e2001164 (2017).

20. Ranjha, L., Anand, R. & Cejka, P. The Saccharomyces cerevisiaeMlh1-Mlh3 Heterodimer Is an Endonuclease That Preferentially Binds to Holliday Junctions. J. Biol. Chem. 289, 5674–5686 (2014).

21. Rogacheva, M. V. et al. Mlh1-Mlh3, A Meiotic Crossover and DNA Mismatch Repair Factor, is a Msh2-Msh3-Stimulated Endonuclease. J. Biol. Chem. (2014). doi:10.1074/jbc.M113.534644

22. de los Santos, T. et al. The Mus81/Mms4 endonuclease acts independently of double-Holliday junction resolution to promote a distinct subset of crossovers during meiosis in budding yeast. Genetics 164, 81–94 (2003).

23. Stahl, F. W. et al. Does crossover interference count in Saccharomyces cerevisiae? Genetics 168, 35–48 (2004).

24. Zalevsky, J., MacQueen, A. J., Duffy, J. B., Kemphues, K. J. & Villeneuve, A. M. Crossing over during Caenorhabditis elegans meiosis requires a conserved MutS-based pathway that is partially dispensable in budding yeast. Genetics 153, 1271–1283 (1999).

25. De Muyt, A. et al. BLM Helicase Ortholog Sgs1 Is a Central Regulator of Meiotic Recombination Intermediate Metabolism. Mol. Cell 46, 43–53 (2012).

26. Matos, J., Blanco, M. G., Maslen, S., Skehel, J. M. & West, S. C. Regulatory control of the resolution of DNA recombination intermediates during meiosis and mitosis. Cell 147, 158–172 (2011).

27. Oke, A., Anderson, C. M., Yam, P. & Fung, J. C. Controlling meiotic recombinational repair - specifying the roles of ZMMs, Sgs1 and Mus81/Mms4 in crossover formation. PLoS Genet. 10, e1004690 (2014).

28. Hunter, N. & Kleckner, N. The single-end invasion: an asymmetric intermediate at the double-strand break to double-holliday junction transition of meiotic recombination. Cell 106, 59–70 (2001).

29. Allers, T. & Lichten, M. Differential timing and control of noncrossover and crossover recombination during meiosis. Cell 106, 47–57 (2001).

30. Börner, G. V., Kleckner, N. & Hunter, N. Crossover/noncrossover differentiation, synaptonemal complex formation, and regulatory surveillance at the leptotene/zygotene transition of meiosis. Cell 117, 29–45 (2004).

31. Alani, E., Reenan, R. A. & Kolodner, R. D. Interaction between mismatch repair and genetic recombination in Saccharomyces cerevisiae. Genetics 137, 19–39 (1994).

32. Nag, D. K., White, M. A. & Petes, T. D. Palindromic sequences in heteroduplex DNA inhibit mismatch repair in yeast. Nature 340, 318–320 (1989).

33. Merker, J. D., Dominska, M. & Petes, T. D. Patterns of heteroduplex formation associated with the initiation of meiotic recombination in the yeast Saccharomyces cerevisiae. Genetics 165, 47–63 (2003).

34. Jessop, L., Allers, T. & Lichten, M. Infrequent co-conversion of markers flanking a meiotic recombination initiation site in Saccharomyces cerevisiae. Genetics 169, 1353–1367 (2005).

35. Allers, T. & Lichten, M. Intermediates of yeast meiotic recombination contain heteroduplex DNA. Mol. Cell 8, 225–231 (2001).

36. Hoffmann, E. R., Eriksson, E., Herbert, B. J. & Borts, R. H. MLH1 and MSH2 promote the symmetry of double-strand break repair events at the HIS4 hotspot in Saccharomyces cerevisiae. Genetics 169, 1291–1303 (2005).

37. Foss, H. M., Hillers, K. J. & Stahl, F. W. The conversion gradient at HIS4 of Saccharomyces cerevisiae. II. A role for mismatch repair directed by biased resolution of the recombinational intermediate. Genetics 153, 573–583 (1999).

38. Pan, J. et al. A Hierarchical Combination of Factors Shapes the Genome-wide Topography of Yeast Meiotic Recombination Initiation. Cell 144, 719–731 (2011).

39. Lam, I. & Keeney, S. Nonparadoxical evolutionary stability of the recombination initiation landscape in yeast. Science 350, 932–937 (2015).

40. Duroc, Y. et al. Concerted action of the MutLβ heterodimer and Mer3 helicase regulates the global extent of meiotic gene conversion. eLife 6, (2017).

41. Smith, C. E., Llorente, B. & Symington, L. S. Template switching during break-induced replication. Nature 447, 102–105 (2007).

42. Llorente, B., Smith, C. E. & Symington, L. S. Break-induced replication: what is it and what is it for? Cell Cycle 7, 859–864 (2008).

43. Schwacha, A. & Kleckner, N. Identification of joint molecules that form frequently between homologs but rarely between sister chromatids during yeast meiosis. Cell 76, 51–63 (1994).

44. Kim, K. P. et al. Sister cohesion and structural axis components mediate homolog bias of meiotic recombination. Cell 143, 924–937 (2010).

45. Chakraborty, U. & Alani, E. Understanding how mismatch repair proteins participate in the repair/anti-recombination decision. FEMS Yeast Research 16, fow071 (2016).

46. Langston, L. D. & O’Donnell, M. DNA polymerase delta is highly processive with proliferating cell nuclear antigen and undergoes collision release upon completing DNA. J. Biol. Chem. 283, 29522–29531 (2008).

47. Cole, F. et al. Mouse tetrad analysis provides insights into recombination mechanisms and hotspot evolutionary dynamics. Nature Publishing Group 46, 1072–1080 (2014).

48. Lange, J. et al. The Landscape of Mouse Meiotic Double-Strand Break Formation, Processing, and Repair. Cell 1–21 (2016). doi:10.1016/j.cell.2016.09.035

49. Abdullah, M. F. F., Hoffmann, E. R., Cotton, V. E. & Borts, R. H. A role for the MutL homologue *MLH2* in controlling heteroduplex formation and in regulating between two different crossover pathways in budding yeast. Cytogenet. Genome Res. 107, 180–190 (2004).

50. Al-Sweel, N. et al. mlh3 separation of function and endonuclease defective mutants display an unexpected effect on meiotic recombination outcomes. doi:10.1101/108498

51. Claeys Bouuaert, C. & Keeney, S. Distinct DNA-binding surfaces in the ATPase and linker domains of MutLY determine its substrate specificities and exert separable functions in meiotic recombination and mismatch repair. PLoS Genet. 13, e1006722 (2017).

52. Wang, T. F., Kleckner, N. & Hunter, N. Functional specificity of MutL homologs in yeast: evidence for three Mlh1-based heterocomplexes with distinct roles during meiosis in recombination and mismatch correction. Proc. Natl. Acad. Sci. U.S.A. 96, 13914–13919 (1999).

53. Thacker, D., Mohibullah, N., Zhu, X. & Keeney, S. Homologue engagement controls meiotic DNA break number and distribution. Nature 510, 241–246 (2014).

54. Bachrati, C. Z. Mobile D-loops are a preferred substrate for the Bloom’s syndrome helicase. Nucleic Acids Res. 34, 2269–2279 (2006).

55. Ira, G., Malkova, A., Liberi, G., Foiani, M. & Haber, J. E. Srs2 and Sgs1-Top3 suppress crossovers during double-strand break repair in yeast. Cell 115, 401–411 (2003).

56. Wu, L. & Hickson, I. D. The Bloom’s syndrome helicase suppresses crossing over during homologous recombination. Nature 426, 870–874 (2003).

57. Mitchel, K., Lehner, K. & Jinks-Robertson, S. Heteroduplex DNA Position Defines the Roles of the Sgs1, Srs2, and Mph1 Helicases in Promoting Distinct Recombination Outcomes. PLoS Genet. 9, e1003340 (2013).

58. Oh, S. D., Lao, J. P., Taylor, A. F., Smith, G. R. & Hunter, N. RecQ Helicase, Sgs1, and XPF Family Endonuclease, Mus81-Mms4, Resolve Aberrant Joint Molecules during Meiotic Recombination. Mol. Cell 31, 324–336 (2008).

59. Kaur, H., De Muyt, A. & Lichten, M. Top3-Rmi1 DNA Single-Strand Decatenase Is Integral to the Formation and Resolution of Meiotic Recombination Intermediates. Mol. Cell 57, 583–594 (2015).

60. Keelagher, R. E., Cotton, V. E., Goldman, A. S. H. & Borts, R. H. Separable roles for Exonuclease I in meiotic DNA double-strand break repair. DNA Repair (Amst.) 10, 126–137 (2011).

61. Guo, X. & Jinks-Robertson, S. Roles of exonucleases and translesion synthesis DNA polymerases during mitotic gap repair in yeast. DNA Repair (Amst.) 12, 1024–1030 (2013).

62. de Massy, B. Distribution of meiotic recombination sites. Trends in Genetics 19, 514–522 (2003).

63. Baker, M. D. & Birmingham, E. C. Evidence for Biased Holliday Junction Cleavage and Mismatch Repair Directed by Junction Cuts during DoubleStrand-Break Repair in Mammalian Cells. Mol. Cell. Biol. 21, 3425–3435 (2001).

64. Mitchel, K., Zhang, H., Welz-Voegele, C. & Jinks-Robertson, S. Molecular Structures of Crossover and Noncrossover Intermediates during Gap Repair in Yeast: Implications for Recombination. Mol. Cell 38, 211–222 (2010).

65. Yin, Y., Dominska, M., Yim, E. & Petes, T. D. High-resolution mapping of heteroduplex DNA formed during UV-induced and spontaneous mitotic recombination events in yeast. eLife 6, (2017).

66. Crown, K. N., McMahan, S. & Sekelsky, J. Eliminating Both Canonical and Short-Patch Mismatch Repair in Drosophila melanogaster Suggests a New Meiotic Recombination Model. PLoS Genet. 10, e1004583 (2014).

67. Bell, L. R. & Byers, B. Homologous association of chromosomal DNA during yeast meiosis. Cold Spring Harbor Symposia on Quantitative Biology 47 Pt 2, 829–840 (1983).

68. Osman, F., Dixon, J., Doe, C. L. & Whitby, M. C. Generating crossovers by resolution of nicked Holliday junctions: a role for Mus81-Eme1 in meiosis. Mol. Cell 12, 761–774 (2003).

69. Dehé, P.-M. & Gaillard, P.-H. L. Control of structure-specific endonucleases to maintain genome stability. Nat. Rev. Mol. Cell Biol. 18, 315–330 (2017).

70. Zhang, L., Kleckner, N. E., Storlazzi, A. & Kim, K. P. Meiotic double-strand breaks occur once per pair of (sister) chromatids and, via Mec1/ATR and Tel1/ATM, once per quartet of chromatids. Proc. Natl. Acad. Sci. U.S.A. 108, 20036–20041 (2011).

71. Vakirlis, N. et al. Reconstruction of ancestral chromosome architecture and gene repertoire reveals principles of genome evolution in a model yeast genus. Genome Res. (2016). doi:10.1101/gr.204420.116

72. Hoffmann, E. R. & Borts, R. H. Trans events associated with crossovers are revealed in the absence of mismatch repair genes in Saccharomyces cerevisiae. Genetics 169, 1305–1310 (2005).

